# Verify Hub Genes of expression profile in aortic dissection

**DOI:** 10.1101/627125

**Authors:** Weitie Wang, Qing Liu, Yong Wang, Hulin Piao, Bo Li, Zhicheng Zhu, Dan Li, Tiance Wang, Rihao Xu, Kexiang Liu

## Abstract

**Background:** To assess the mRNAs expression profile and explore the hub mRNAs and potential molecular mechanisms in the pathogenesis of human thoracic aortic dissection (TAD). Methodology: mRNA microarray expression signatures of TAD tissues (n=6) and no TAD tissues (NT;n=6) were analyzed by Arraystar human mRNAs microarray. Real-time PCR (qRT-PCR) were used to validate the result of mRNAs microarray. Bioinformatic tools including gene ontology, and Kyoto Encyclopedia of Genes and Genomes pathway analysis were utilized. The protein-protein interaction networks were constructed based on data from the STRING database. Molecular Complex Detection (MCODE) and cytohubba analysis were used to infer the most hug gene and pathways. Results: The top 10 hub genes CDK1, CDC20, CCNB2, CCNB1, MAD2L1, AURKA, C3AR1, NCAPG,CXCL12 and ASPM were identified from the PPI network. Module analysis revealed that TAD was associated with cell cycle, oocyte meiosis, p53 signaling pathway, progesterone-mediated oocyte maturation. The qRT-PCR result showed that the expression of all hug genes was significantly increased in TAD samples (p < 0.05). Conclusions: These candidate genes could be used as potential diagnostic biomarkers and therapeutic targets of TAD.

**Author summary:** Many basic characteristics underlying the establishment of aortic dissection have not been studied in detail. The presented work sought to understand the pathogenesis of human thoracic aortic dissection by employing bioinformatic tools to explore the hub mRNAs and potential molecular mechanisms of thoracic aortic dissection. Many pathway were thought to have relevant with this disease, but the most important pathway was not define. We used bio-mathematical analysis to explore the potential functions in thoracic aortic dissection and identified the hub genes and explored the intrinsic molecular mechanisms involved in thoracic aortic dissection between two microarray analysis. Finally, we indentified the cell cycle maybe the key pathway in thoracic aortic dissection.

## Introduction

Thoracic aortic dissection (TAD) is a common and a life-threatening aortic disease ^1^. Despite improvements in medical therapy and surgical or endovascular techniques in recent years, TAD still remains a high morbidity and mortality rate ^2^. Owing to the poor results of the existing treatment methods, necessary understanding of the molecular mechanism maybe provided new insights into therapeutic targets for TAD. Many reports show the degradation of extracellular matrix (ECM) and depletion of vascular smooth muscle cell (VSMC) of the aortic wall are the main histopathological findings ^3-5^. However, it is remains unclear of the key molecular mechanism of TAD pathogenesis.

In recent years, mRNAs have been reported to participate in the regulation of pathophysiological conditions and have been proved to involve in the progression of cardiovascular disease ^6^. Nowadays, researches have focused on the Genomewide Association Studies (GWAS) ^7^, which could find relevant genetic variants that may be used as potential biomarkers for diagnosis and targeted therapy. Although high-throughput sequencing technology have provided many diverse expressed genes, different expression profile in TAD provided various results and no reliable results have been identified up to now ^8-10^. More samples and advanced bio-informatics methods should be used in further study.

In the present study, we used mRNA microarray to acquire differential expression profiles in human TAD tissues and CAD tissues. Subsequently, bio-mathematical analysis were used to explore the potential functions in TAD and identified the hub genes and explore the intrinsic molecular mechanisms involved in TAD. Thus, these data would provide a foundation for new biomarkers and therapeutic targets for human TAD.

## RESULTS

### Differential expression profiles of mRNAs in TAD and NT group

Volcano plots revealed that mRNAs were differentially expressed in human TAD aortic tissues through microarray technology (Figure 1). A total of 2834 mRNAs were differentially expressed in TAD group compared with NT group. In total, 1928 mRNAs were up-regulated, and 906 mRNAs were down-regulated (fold change >2.0, P value < 0.05). The top ten up- and down-regulated mRNAs are listed in Table 3.

**Figure 1.**
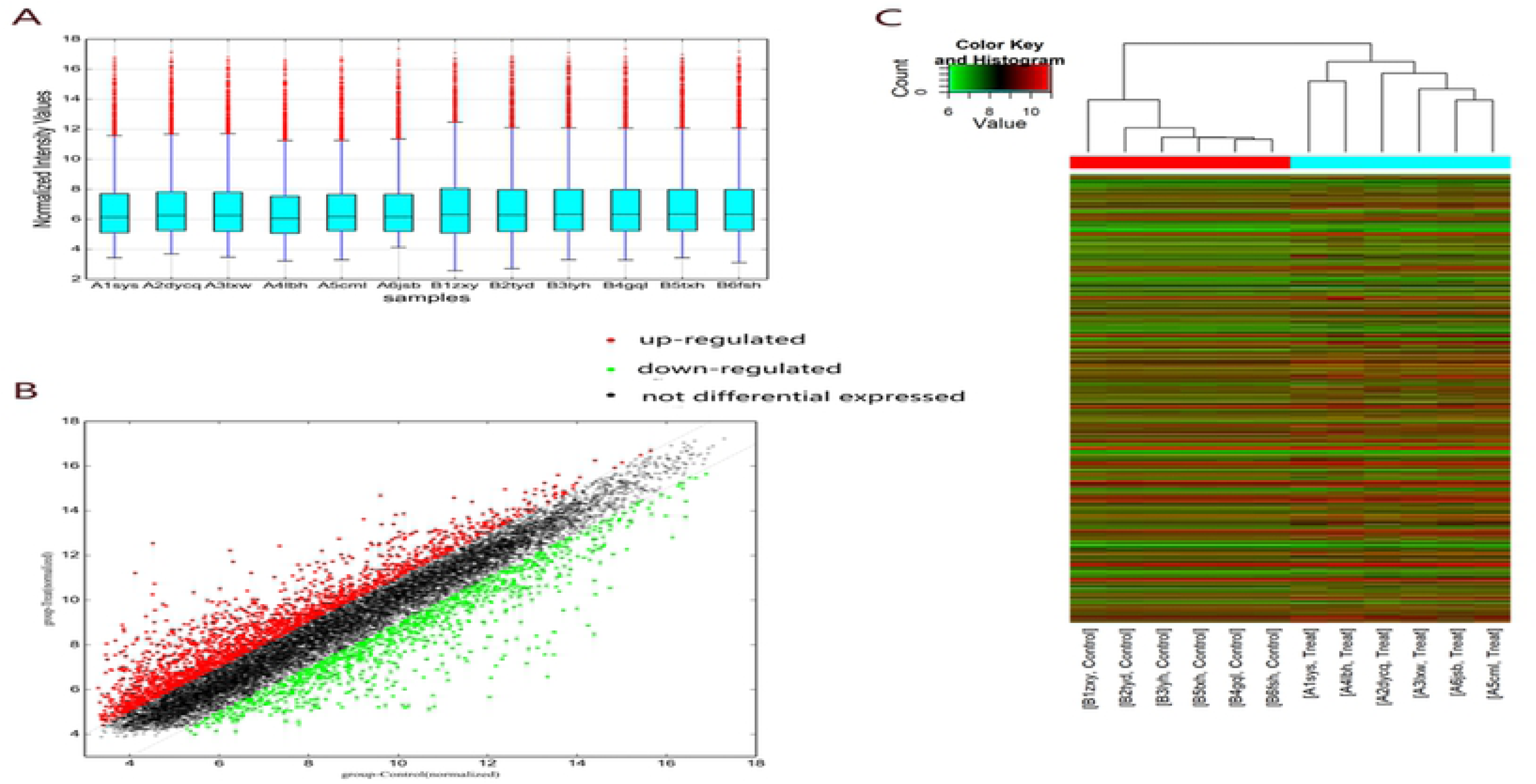
Comparison of mRNA expression profile between the TAD samples and NT samples. Comparison of mRNA expression profile between the TAD samples and NT samples. (A) The box plot is a convenient method to quickly compare the distribution of mRNAs. After normalization, the distributions of log2 ratios among the tested samples are almost similar. (B) The scatterplot is a visualization method that is useful for assessing the variation between the TAD and control tissues compared by microarrays. The values of X and Y axes in the scatterplot are averaged normalized values in each group (log 2 scaled). The green plot are decrease and red plot are upgrade different gene.(C)diferentially expressed genes can be effectively divided into TAD and NT groups. Red indicates that the gene that is upregulated and green represents down-regulated genes.

### GO Functional Enrichment Analysis

To know the functions of all DEGs, we used DAVID online tool, the DEGs functions of GO function enrichment were divided into three groups including BP, CC, and MF (Figure 2). As shown in the Figure 2 and Table 4, in the biological processes group, the down-DEGs are mainly enriched in cardiac muscle tissue development, muscle tissue development, muscle structure development, developmental process, and single-multicellular organism process and the up-DEGs are mainly enriched inimmune response, response to stress, defense response, immune system process and innate immune response. In the cellular component group, the down-DEGs are mainly enriched in contractile fiber, myofibril, sarcomere, contractile fiber part and cell-substrate junction and the up-DEGs are mainly enriched in cytoplasm, cytoplasmic part, intracellular organelle part, organelle part and membrane-bounded organelle. And in the molecular function group, the down-DEGs are mainly enriched in protein phosphatase regulator activity, phosphatase regulator activity, cytoskeletal protein binding, structural constituent of muscle and actin binding and the up-DEGs are mainly enriched in catalytic activity, protein binding, single-stranded DNA-dependent ATPase activity, ATP binding and adenyl ribonucleotide binding.

**Figure 2.**
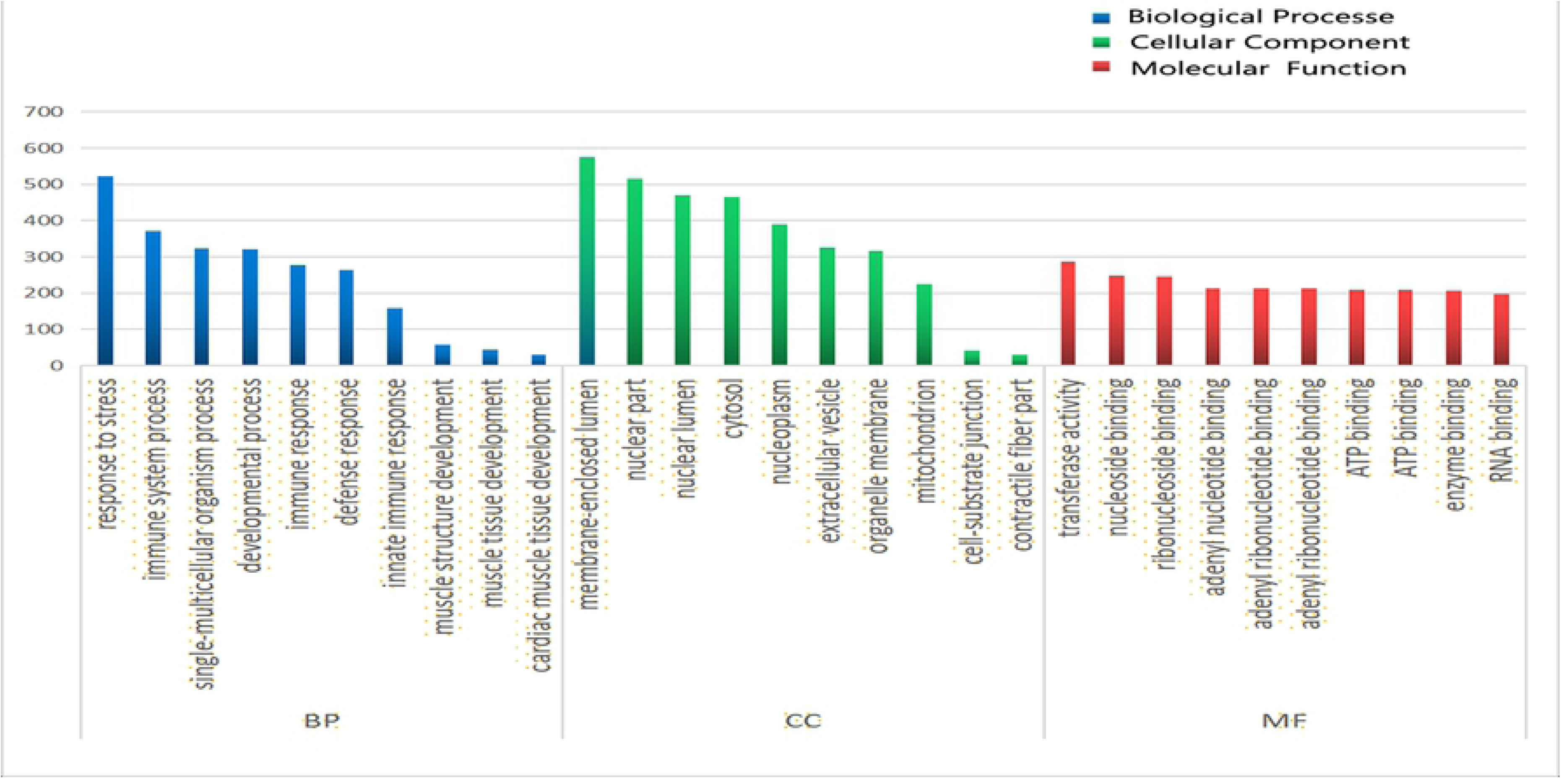
Gene Ontology analysis classified the differentially expressed genes into 3 groups. molecular function, biological process, and cellular component.

### Signaling Pathway Analysis

After the pathway enrichment analysis, down regulated genes were mainly enriched in dilated cardiomyopathy, arrhythmogenic right ventricular cardiomyopathy, hypertrophic cardiomyopathy, adrenergic signaling in cardiomyocytes and vascular smooth muscle contraction. And up regulated genes were mainly enriched in DNA replication, phagosome, cell cycle, staphylococcus aureus infection and lysosome (Figure 3,4).

**Figure 3.**
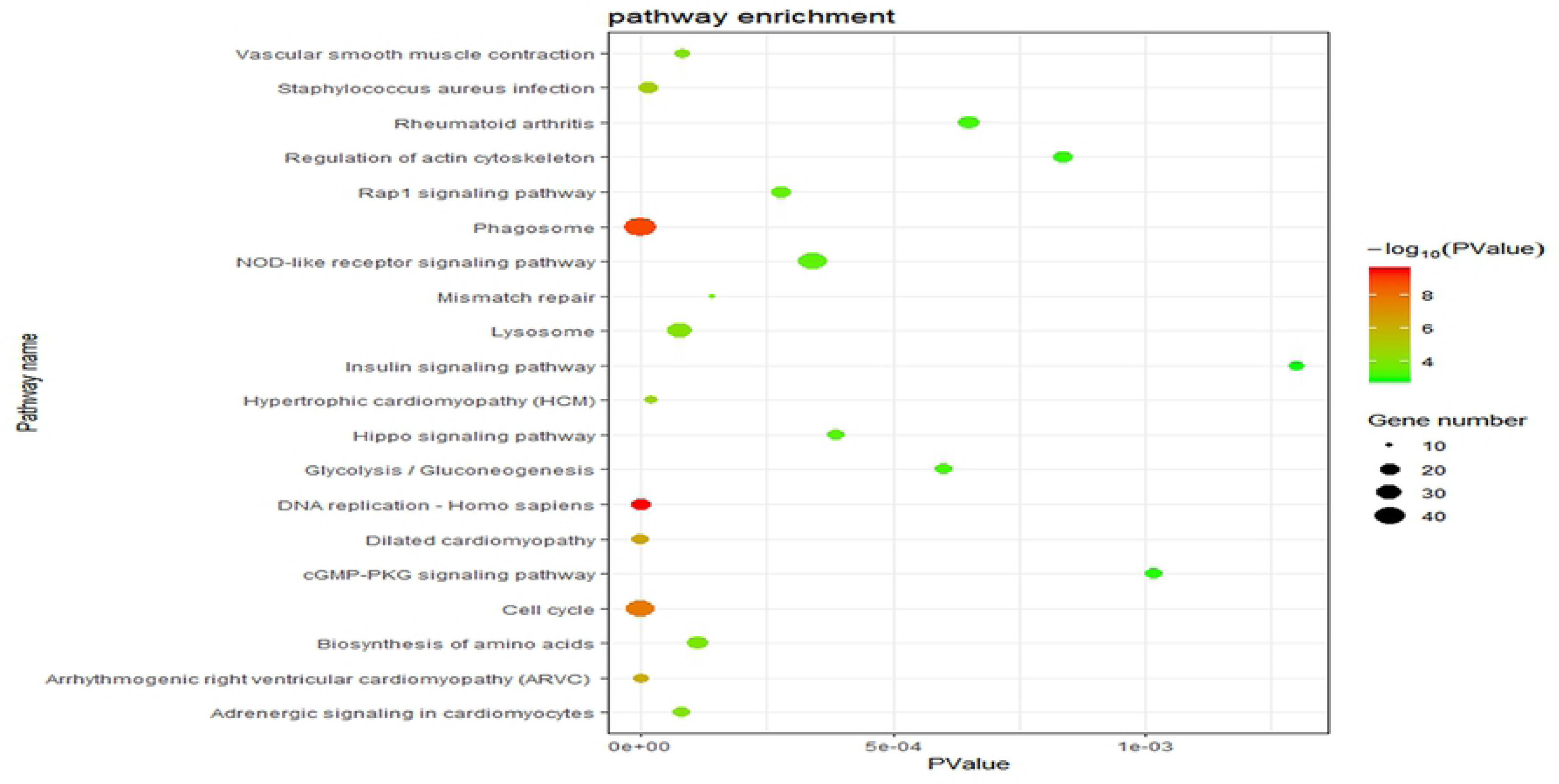
Kyoto Encyclopedia of Genes and Genomes enrichment analysis of the pathways. The gradual color represents the P value; the size of the black spots represents the gene number.

**Figure 4.**
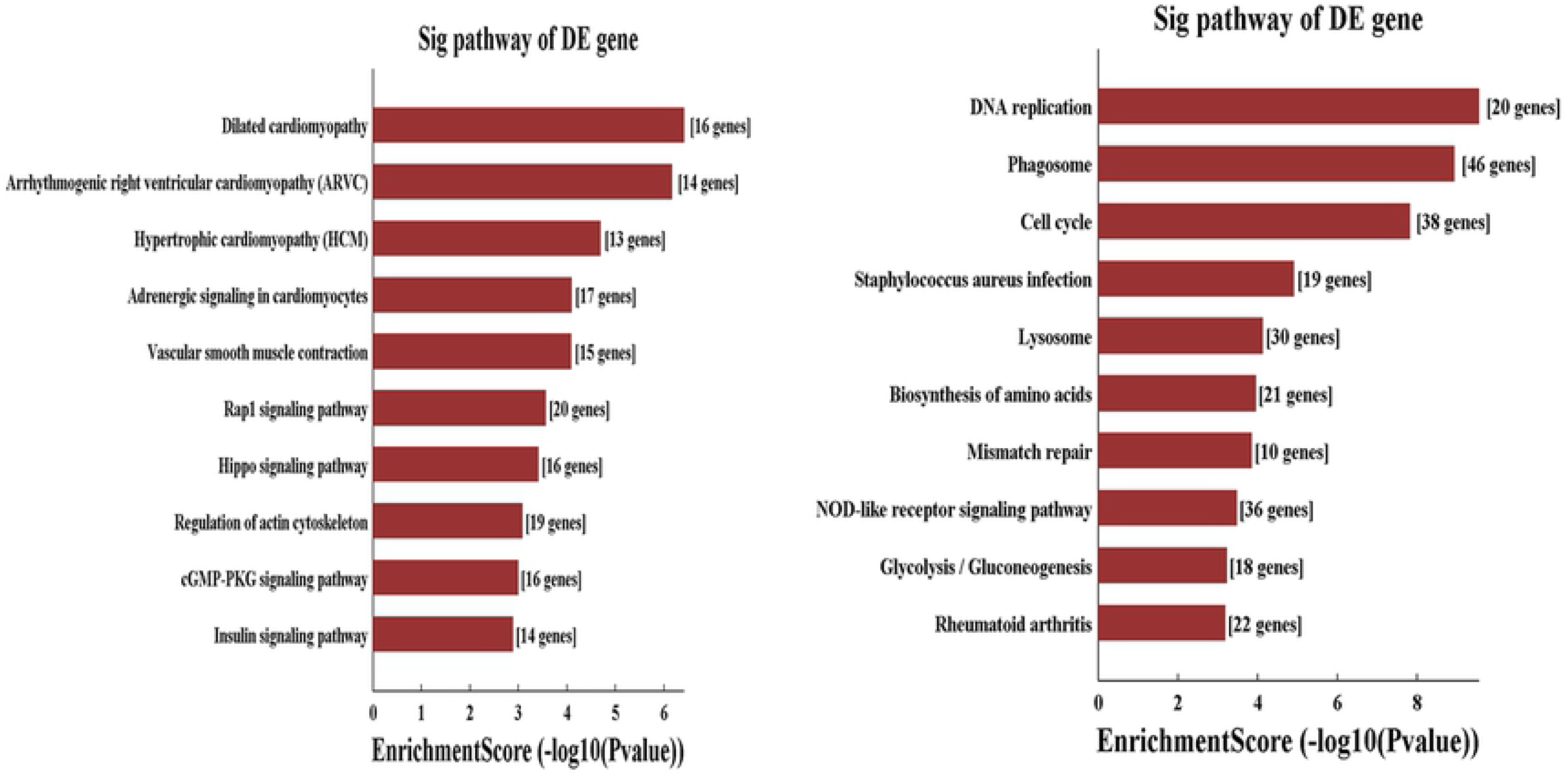
Signifcantly up-regulated and down-regulated pathways.

### PPI Network and Modular Analysis

All the DEGs (fold change >4.0) were analysis using STRING database. Then we put these data into Cytoscape software to construct a PPI network which containing 433 nodes and 348 edges (Figure 5). In these DEGs, 10 hub genes including CDK1, CDC20, CCNB2, CCNB1, MAD2L1, AURKA, C3AR1, NCAPG,CXCL12, ASPM after calculating. In these 10 hub genes, CDK1 present with the highest degree (degree = 44). The Cytoscape plugin MCODE shows the top three module with 15.429,14.857 and 10.000 score respectively (Figure 6). Then the genes in these three module were performed with functional enrichment analyses. Pathway enrichment analysis showed that Module 1 is mainly relevant with cell cycle oocyte meiosis, p53 signaling pathway, progesterone-mediated oocyte maturation. Module 2 is mainly associated with chemokine signaling pathway, cytokine-cytokine receptor interaction, rheumatoid arthritis, NF-kappa B signaling pathway, neuroactive ligand-receptor interaction. Module 3 is mainly associated with leukocyte transendothelial migration. Validation by qRT-PCR of differentially expressed mRNAs

**Figure 5.**
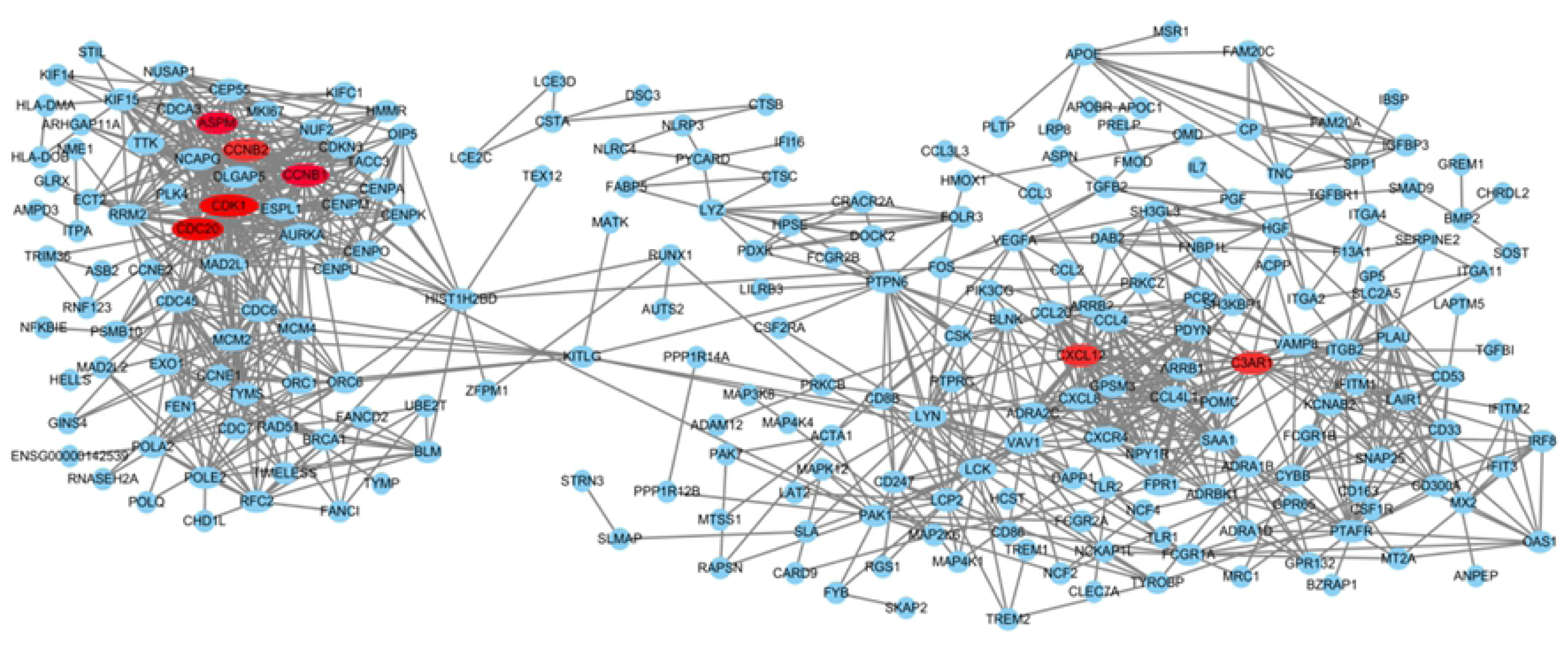
PPI network constructed with the differentially expressed genes. PPI network constructed with the differentially expressed genes. Red nodes represent hub GENE analysis by cytohubba.

**Figure 6:**
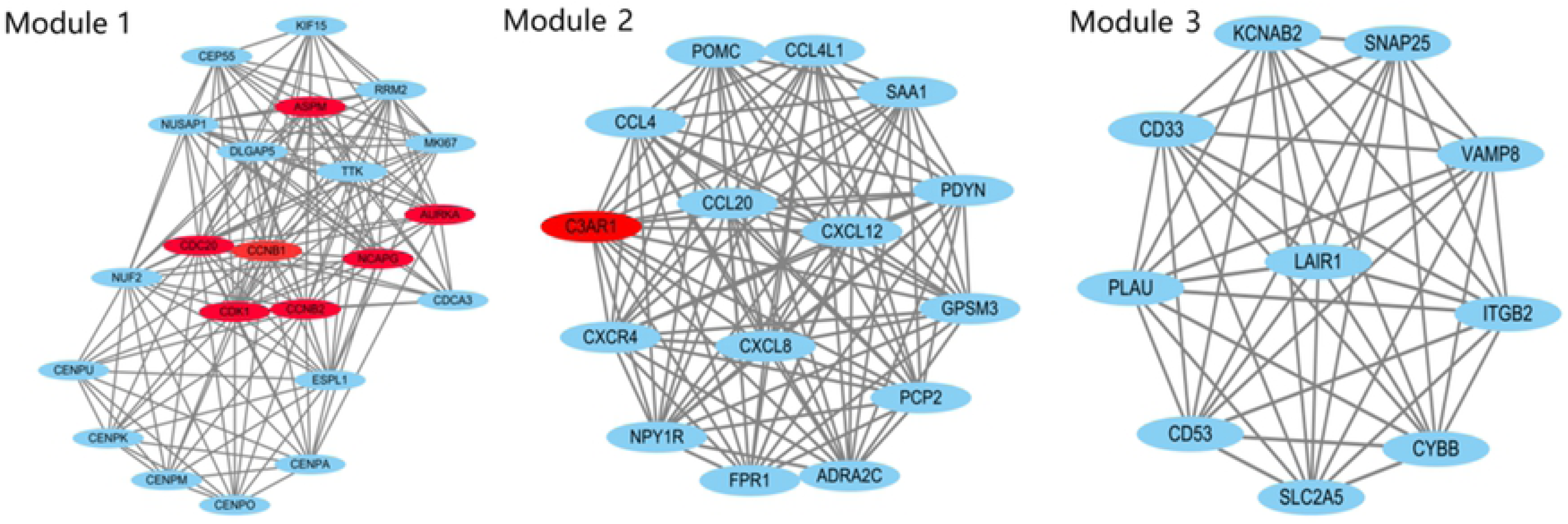
The three most significance modules. Red nodes represent hug GENE analysis by cytohubba.

To validate microarray results, the expression levels of top 10 hub genes were determined in ascending aortic samples of thoracic aortic dissection and no thoracic aortic dissection using qRT-PCR. The verification result showed that the expression levels of the 10 hub genes were significantly increased in thoracic aortic dissection samples (p < 0.05) (Figure 7). All validations are consistent with the microarray data and analytical results in this study.

**Figure 7.**
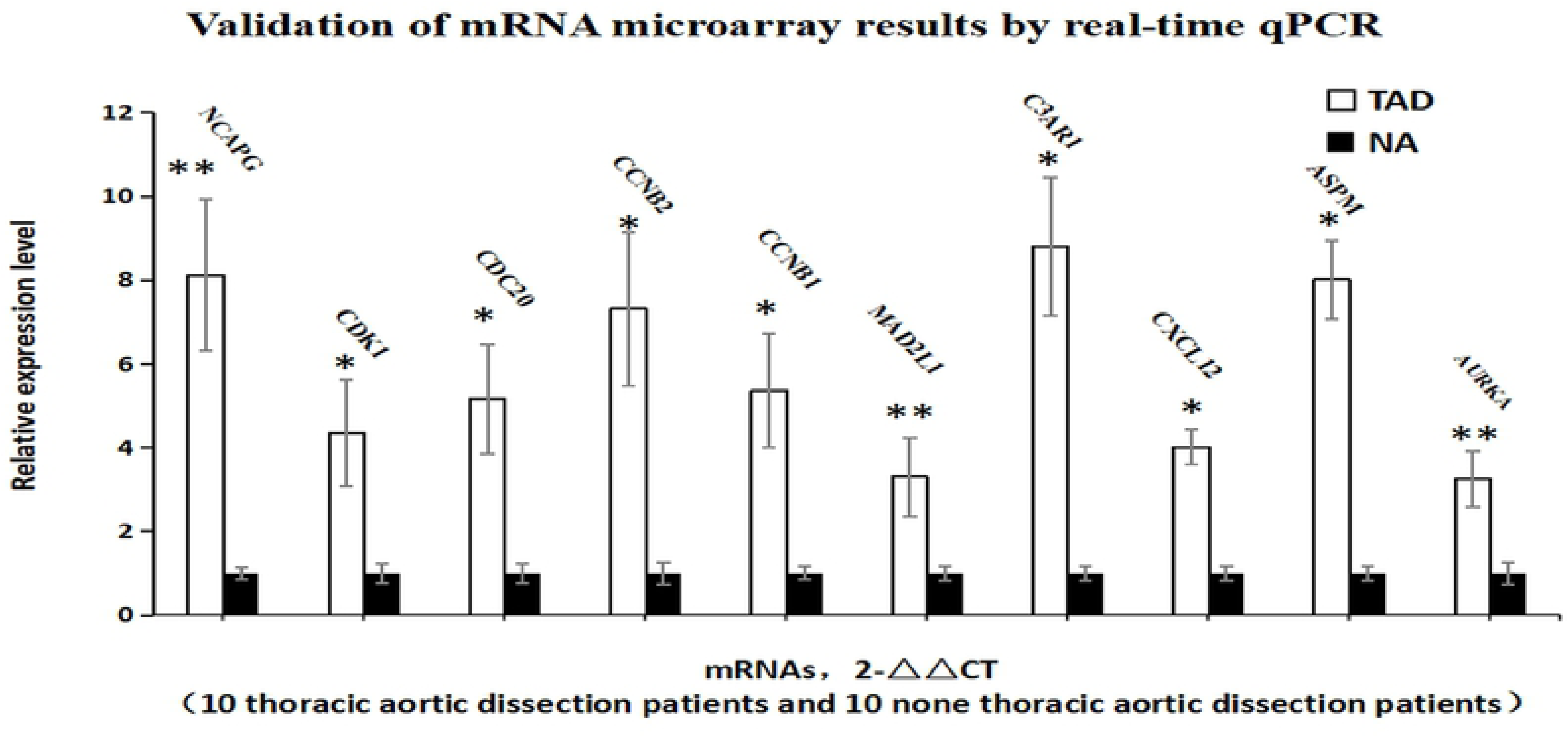
Validation of the mRNA microarray results by real-time qPCR. mRNA microarray result was verified by real-time qPCR between the TAD group (n = 10) and the NT group (n = 10). All samples were normalized to the expression of GAPDH, and the relative expression levels of each gene were analyzed using the 2-△△Ct method. *P < 0.05, **P < 0.01.

## Disscussion

TAD is a life-threatening event that carries a high mortality rate ^1^. Patients always complain with acute chest and back pain and are often misdiagnosed as acute myocardial infarction ^15^. Surgical treatment seems to the best method of this disaster disease ^16^. However, the traditional surgical treatment of TAD is very complex and time-consuming with high mortality and poor prognosis. Therefore it is necessary to research of the new biomarkers and therapeutic targets. Up to now, the molecular mechanism of this severe disease remains unclear. TAD such as Marfan syndrome and Ehlers-Danlos syndrome had been confirmed with deficiency of the glycoprotein ^17^ and abnormal type-III precollagen ^18^ respectively. However, most patients with TAD do not exhibit such explicit syndromes. These TAD patients always complain with hypertension, atherosclerosis and trauma ^15^. In addition, lots of studies had pointed that degradation of ECM and depletion of VSMC play an important role in un-heredity TAD ^19^. So studies are primarily focused on the protein-coding genes of ECM ^20^ and VSMC such as COL3A1, FBN1, LOX, FLNA, ACTA2, MYH11. These protein-coding genes play role in pathological processes of TAD. However, the key pathogenesis of the TAD has not been confirmed up to now.

Modern molecular biology believe that the disease was induced by the changing of various gene expression profiles of tissues or cells. Collection, summary and analysis of the large numbers of these gene can help us to understand the mechanism of the diseases. High-throughput sequencing technology make the identity of the changing gene in tissue available and have been widely used to predict potential targets gene. Up to now, there are many studies have been performed and thousands of differentially expressed genes had been screened. However, different gene profile are various with each study ^8-10^. So more further analysis should be carried out and hub genes and pathway should be identify.

This study identify 2834 mRNAs DEGs which included 1928 upregulated DEGs and 906 downregulated DEGs (fold change >2.0, P value < 0.05). Like other studies ^8-10^, we classified into three groups including BP,CC, PF by GO terms. GO functional enrichment analysis showed that immune response, response to stress, defense response and immune system process, which was consistent with other studies. Inflammatory and immunological may both probably lead to the degree of vascular damage of aortic wall ^21-23^. Reports had presented that inflammatory mechanisms participated in medial degeneration of aortic dissection tissue. The macrophages and activated T lymphocytes were also found in the dissection tissue. T helper 2 response was reported to have relevant with the growth of aneurysms ^24-25^. hyper-expression IL-6 and IL-8 in aortic dissection was also present that immunologic pathways were critical in the aortic wall damage ^26^. In addition, neutrophils, CD8+, CD28–,IL-6, TNF-α, IL-8, and MCP-1 suggestted that cytotoxic and innate cells were mainly relevant with the pathogenesis of TAD ^27^. T helper lymphocytes and activated macrophages, mast cells, T and B lymphocytes that immune system process and inflammatory pathways were involved in the weakening of the aortic wall.

Furthermore, the enriched KEGG pathways of downregulated genes were mainly enriched in dilated cardiomyopathy, arrhythmogenic right ventricular cardiomyopathy, hypertrophic cardiomyopathy, adrenergic signaling in cardiomyocytes and vascular smooth muscle contraction. VSMC was the major cell in aortic media. The function of VSMC such as proliferation and migration acted critical biomechanical properties of the aortic wall. It was affluence by many factor including lymphocytes induce apoptosis of VSMCs and synthesis of MMPs. So many studies aim to research the function of VSMC. Talin-1 had been reported to main exist in aortic media and it was significant downregulation in AD aortic tissue. Further study confirmed that Talin-1 regulating VSMC proliferation and migration and finally caused of pathologic vascular remodeling to change vascular media structure and function and lead to AD ^28^. Many gene such as YAP1, Sirtuin-1, PCSK9, polycystin-1, brahma-related gene 1 had also been reported to associated with the pathophysiologic processes of aortic dissection through influence proliferation and migration of VSMC ^29-31^.

Like previous high-throughput sequencing studies, we also got many relevant pathways through analysising, but we can not sure which one is the most relevant mechanism about TAD and the most hug gene about AD is also not sure. So we carry out some further analysis such as PPI Network and Modular Analysis including STRING database, cytohubba analysis and MCODE analysis which is mainly widely used in infer the most hug gene and pathways ^32^. In our study, we choose gene which is different express up to 4 fold in order to reduce the scope. The cytohubba analysis show that the top ten hug genes are CDK1, CDC20, CCNB2, CCNB1, MAD2L1, AURKA, C3AR1, NCAPG,CXCL12, ASPM. These gene shows the cell cycle seem to play an important role in TAD. So the Cytoscape plugin MCODE was used to analysis from another aspect. After calculating, the top module including 22 nodes and 162 edges. We choose the 22 nodes for functional enrichment analyses. And the cell cycle was the most relevant pathway and also include 7 hug genes which previous forcast. So we aim to analysis the hug gene CDK1 and cell cycle in TAD.

CDK1 is the founding member of the CDK family in human cells ^33^. It can conserve across all leukocytes and is the only essential cell cycle CDK in human cells. Study reported that this important member of CDK family is required for successful completion of M-phase. In addition, the conserved nature and remodelled function make this molecular more special. In cell cycle, CDK1 is always act it function combine with cyclin A and cyclin B. So CDK1 can regulate the cell cycle from GI to S phase because it partner-cyclin A-is first expressed during late G1 where it initially binds to CDK2 and promotes S-phase ^34^. As the cyclin-dependent kinase (CDK) family in association with partner cyclin proteins will mediated the progression of cell cycle, CDK1 inhibition will lead the initiates adhesion remodeling in preparation for entry into mitosis and reveal an intimate link between the cell cycle machinery and cell–ECM adhesion. So the toppest degree hug gene is CDK1 which is mean the cell cycle will act as the key role in TAD. The Cytoscape plugin MCODE result also show the first progosis passway is cell cycle. The two analysis method are consistent and also in accordence with previous studies ^8-10^.

The endothelial cells and SMC are important in maintaining of vascular tone in blood vessel. Previous study confirm that restricting cell cycle of endothelial cell and SMC may lead to disorder of vascular remodeling as an antecedent to the pathological sequelae of cardiovascular disease ^35^. The SMC function changing are two main pathogenesis of TAD. SMC progression and proliferation are critical for the development of TAD. So most studies aim to research the cell cycle of SMC ^36^. Arrests proliferating SMCs in G0/G1 and G2/M-phases and inhibits cyclin B1 expression and cdk1 activity (essential for G2-to-M progression) may protect vascular-proliferative diseases. Other investigations also revealed an important role of VSMCs proliferative in aortic tissue degeneration, which has been believed as the initiation of pathologic remodeling in TAD ^37^. it is obvious that the abnormal proliferation and migration of VSMCs and changing of cell cycle have been proved to the considered as the potentially main cause of pathological vascular remodeling through undermining the vasculature stability and finally lead to vascular disease. many studies aim to research the VSMCs from dissected aorta and indentidy this SMC can proliferated more rapidly than normal VSMCs tissues, and the genes participate in proliferation exhibited an increased expression. In our study, the 10 hug genes are all up expression which show the up regulaition defferent expression will act an important role in TAD which is coninsistance with previous study.

In summary, by means of high-throughput sequencing and data processing as well as qRT-PCR validation, the hub genes including CDK1, CDC20, CCNB2, CCNB1,AURKA,NCAPG and ASPM may have the potential to be used as drug targets and diagnostic markers of TAD. Cell cycle maybe the key pathway in TAD. However, there were still some limitations: normal aortic tissue used as control group maybe more accuracy. Further experimental studies with larger sample size need to confirm cell cycle pathway in TAD.

## Methods

### Tissue collection

This study was conducted in accordance with the Declaration of Helsinki and was approved by the Ethics Committee of the Second Hospital of Jilin University. Ascending aortic specimens near the intimal tear were obtained from TAD patients undergoing surgical repair (n=6) if the informed consent was obtained. Aortas specimens derived from patients without aortic diseases undergoing coronary artery bypass (n=6) graft were obtained as no TAD (NT) group if the informed consent was obtained. General information of all patients was shown in Table 1. There were no significant differences in age, gender, obesity, smoking between the two groups. Patients diagnosed with TAD were confirmed by the computed tomography angiography (CTA) and excluded from any heredity TAD. All the specimens were immediately frozen in liquid nitrogen, and preserved at −80° C for microarray analysis and qRT-PCR or further usage.

**Table 1.** General information of all patients

|  | TAD group | NT group | P value |
| --- | --- | --- | --- |

|  | (n=6) | (n=6) |  |
| --- | --- | --- | --- |
| Age (years old) | 55.61 ± 9.23 | 57.53 ± 10.61 | 0.7450 |
| Male | 3(50.00%) | 3(50.00%) | - |
| Obesity (BMI >25 kg/m2) | 2(33.33%) | 3(50.00%) | 0.5582 |
| Smoking | 2(33.33%) | 4(66.67%) | 0.2482 |
| NYHA class III-IV | 1(16.67%) | 2(33.33%) | 0.5055 |
| Hypertension | 4(66.67%) | 3(50.00%) | 0.5582 |
| Diabetes mellitus | 1(16.67%) | 3(50.00%) | 0.2207 |
| Chronic renal dysfunction | 0 | 0 | - |

TAD: Thoracic aortic dissection; NT: no TAD; NYHA : New York Heart Association

### RNA extraction and quality control

Tissue RNA from TAD and NT group was extracted according to the manufacturer’s instructions by TRIzol reagent (Invitrogen, NY, USA). NanoDrop ND-1000 (Thermo Fisher Scientific, Wilmington, DE, USA) and Agilent 2100 Bioanalyzer (Agilent Technologies) were used to check the integrity and concentration of the RNA samples. All the qualified samples were stored at −80 °C for future experiments.

### Microarray analysis

There are 12 samples (6 TAD and 6 NT) were used for microarray analysis by Arraystar Human mRNA microarray which contained 14596 mRNAs probes. Sample preparation and array hybridization were performed according to the Agilent One-Color Microarray-Based Gene Expression Analysis protocol (Agilent). Briefly, mRNA was purified from total-RNA (RNeasy Mini Kit, Qiagen) after removal of rRNA and transcribed into fluorescent cRNA without 3’ bias along the entire length of the transcripts with a random priming method. Quick-Amp Labeling Kit (Agilent, USA) was used for sample labeling. Then NanoDrop ND-1000 was used to check the specific activity and concentration of the labeled cRNAs. Then hybridization was performed in an Agilent Hybridization Oven. Data normalization and processing was performed using the GeneSpring GX v12.1 software package (Agilent). After quantile normalization of the raw data, mRNAs samples had flags in Present or Marginal (“All Targets Value”) were chosen for further data analysis. The differentially expressed mRNAs were statistical significance with the change in threshold values was >2.0 or < −2.0 fold between the two groups and Benjamini-Hochberg corrected P <0.05.

### Gene Ontology (GO) and Pathway Enrichment Analyses

DAVID (the Database for Annotation, Visualization, and Integrated Discovery) online bio-informatics database is an analysis tools of biological data to integrates and provide information for biological function and protein list ^11^. This tool was used in this study to provide GO enrichment and Kyoto Encyclopedia of Genes and Genomes (KEGG) pathway analysis. GO analysis included categories of cellular component (CC), biological processes (BP) and molecular function (MF). Pathway analysis is a functional analysis that maps genes to KEGG pathways. And gene count >2 and p < 0.05 were set as the cutoff point.

### Integration of Protein-Protein Interaction (PPI) Network Analysis

STRING (https://string-db.org/cgi/input.pl) is an online database resource search tool which can provide analysis of interacting genes including physical and functional associations ^12^. In this study, the STRING online tool was used to construct a PPI network of upregulation and downregulation diferentially expressed genes (DEGs), with a confidence score >0.7 defined as significant. Then the interaction data were typed into the Cytoscape software ^13^ to structure a PPI network. Based on the above data, we used Molecular Complex Detection (MCODE) ^14^, a built-in APP in Cytoscape software, to analyze the interaction relationship of the DEGs encoding proteins and screening hub gene. The parameters of network scoring and cluster finding were set as follows: degree cutoff = 2, node score cutoff = 0.2, k-core = 2, and max depth = 100.

### Quantitative reverse transcription-PCR (qRT-PCR)

Validation and Statistical Analysis. qRT-PCR was used to verify the core genes. Total RNA was reverse-transcribed to cDNA using PrimeScript RT reagent Kit with gDNA Eraser (TaKaRa, Japan) according to the manufacturer’s instructions. Primer 5.0 software (PREMIER Biosoft, Palo Alto, CA, USA) was used to design primers, and a QuantStudio 7 Flex real-time PCR system (Applied Biosystems, Carlsbad, CA, USA) was used. All primers used in this study were listed in Table 2. All samples were normalized to GAPDH. And the relative expression levels of each gene were calculated using 2-ΔΔCt methods.

**Table 2:**
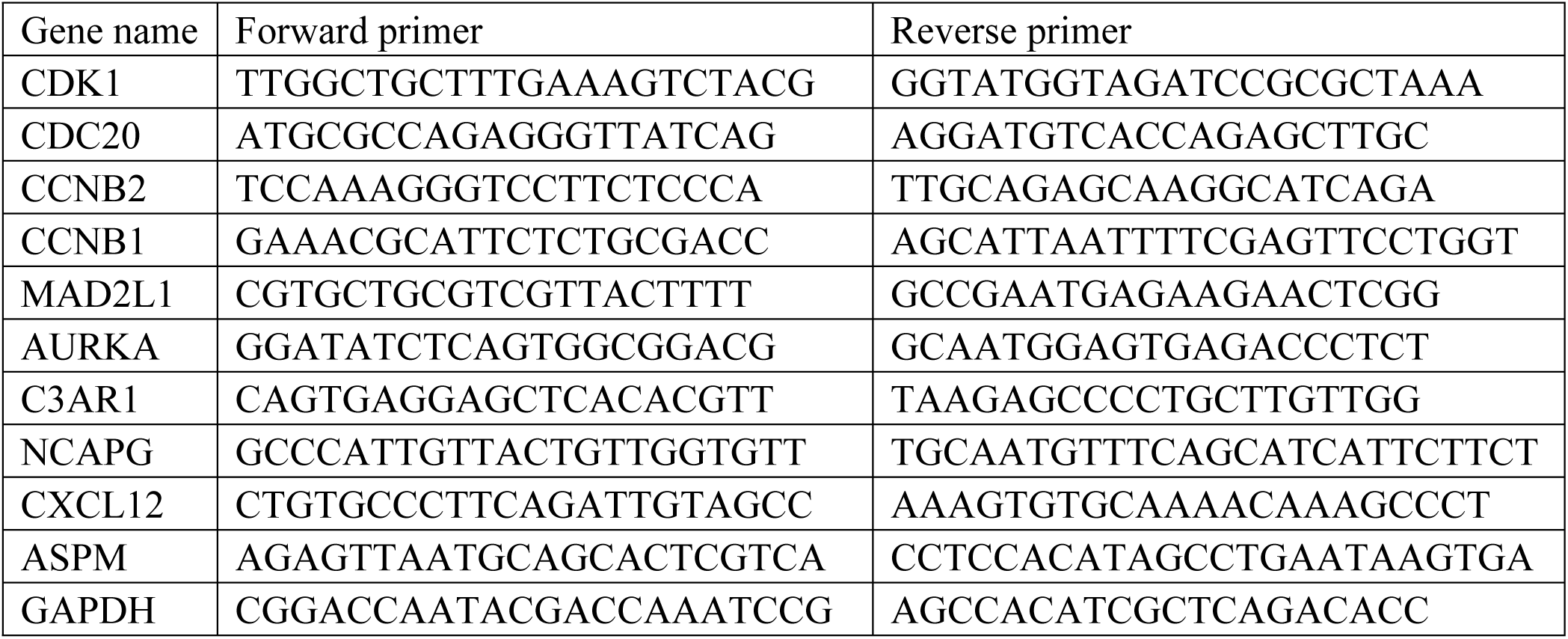
The primers of top 10 hub genes

**Table 3:**
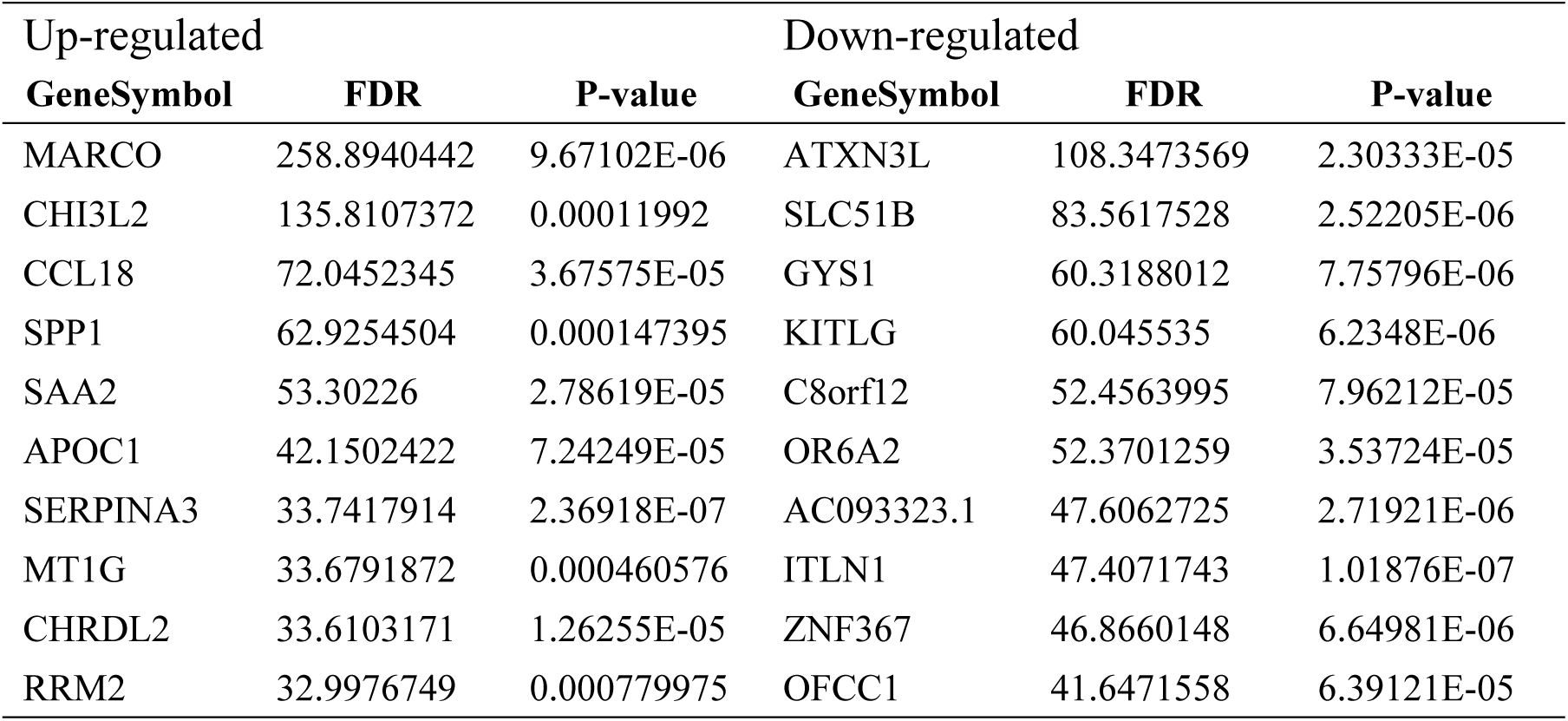
The top ten up- and down-regulated mRNAs

**Table 4.**
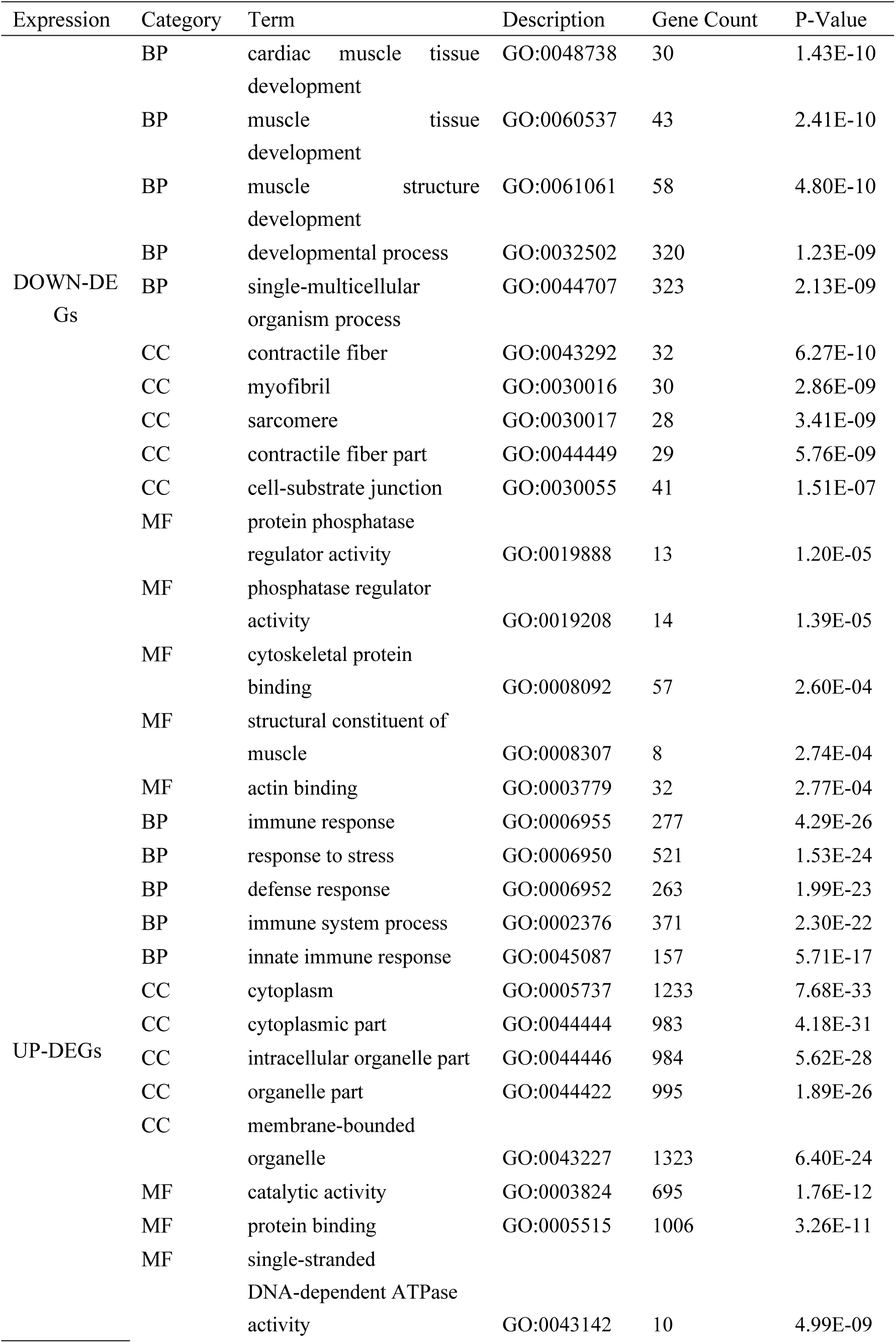
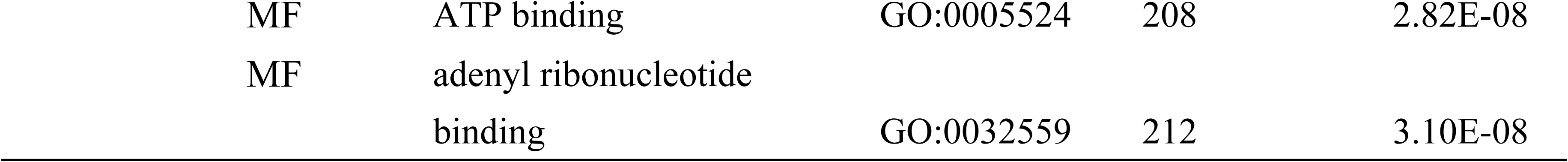
The significant enriched analysis of differentially expressed genes in thoracic aortic dissection

### Statistical analysis

The data were presented as the mean ± standard deviation. Statistical analyses were performed using SPSS version 20.0 (SPSS Inc., Chicago, IL, USA). The raw data were preprocessed by affy package in R software and limma package in R software. Comparisons between groups were performed using unpaired Student’s t-test. Fisher’s exact test was used to evaluate the significance of GO terms and Pathway identifiers enrichment. The false discovery rate (FDR) controlling was used to correct p-value. FDR and P-value less than 0.05 was considered statistically significant.

## Acknowledgments

None

## Contributions

W.T.W, B.L, H.L.P, Q.L and Y.W wrote the paper. Z.C.Z, D.L, T.C.W, R.H.X, and K.X.L checked the References. All authors reviewed the final manuscript.

## Data availability

Data available on request for the Ethics Committee of the Second Hospital of Jilin University.

## Dual publication

The elements in the manuscript have not been published or are under consideration for publication elsewhere.

## Abbreviation

(TAD): thoracic aortic dissection
(ECM): extracellular matrix
(VSMC): vascular smooth muscle cell
(GWAS): Genomewide Association Studies
(NT): no thoracic aortic dissection
(CTA): computed tomography angiography
(GO): Gene Ontology
(KEGG): Kyoto Encyclopedia of Genes and Genomes
(CC): cellular component
(BP): biological processes
(MF): molecular function
(PPI): Protein-Protein Interaction
(DEGs): diferentially expressed genes
(MCODE): Molecular Complex Detection
(qRT-PCR): Quantitative reverse transcription-PCR
(FDR): false discovery rate
(CDK): cyclin-dependent kinase

